# Heading choices of flying *Drosophila* under changing angles of polarized light

**DOI:** 10.1101/665877

**Authors:** Thomas F. Mathejczyk, Mathias F. Wernet

**Affiliations:** Freie Universität Berlin, Fachbereich Biologie, Chemie und Pharmazie Institut für Biologie – Neurobiologie, Königin-Luise Strasse 1-3, 14195 Berlin, Germany

## Abstract

Many navigating insects include the celestial polarization pattern as an additional visual cue to orient their travels. Spontaneous orientation responses of both walking and flying fruit flies (*Drosophila melanogaster*) to linearly polarized light have previously been demonstrated. Using newly designed modular flight arenas consisting entirely of off-the-shelf parts and 3D-printed components we present individual flying flies with a slow and continuous rotational change in the incident angle of linear polarization. Under such open-loop conditions, single flies choose arbitrary headings with respect to the angle of polarized light and show a clear tendency to maintain those chosen headings for several minutes, thereby adjusting their course to the slow rotation of the incident stimulus. Importantly, flies show the tendency to maintain a chosen heading even when two individual test periods under a linearly polarized stimulus are interrupted by an epoch of unpolarized light lasting several minutes. Finally, we show that these behavioral responses are wavelength-specific, existing under polarized UV stimulus while being absent under polarized green light. Taken together, these findings provide further evidence supporting Drosophila’s abilities to use celestial cues for visually guided navigation and course correction.

## Introduction

Like many other animals, insects have developed the ability to efficiently navigate the most complex environments. Over several decades, evidence has accumulated showing that different insect species combine a multitude of visual stimuli in order to take fast and reliable navigational decisions (reviewed in: ^1^). Amongst these cues, the celestial polarization pattern serves as a robust visual stimulus informing the heading choices of many navigating insects ^1–3^. Since Karl von Frisch first described the ability of honeybees to orient their waggle dances using merely a small patch of sky that did not include the sun as a landmark, many insects have also been shown to integrate the directional information provided by the skylight polarization pattern into their repertoire of visual cues ^4^. Importantly, this ability is not restricted to central-place foragers like bees or desert ants that rely on visual cues to find their way back to their hive or nest ^5,6^. For example, both diurnal and nocturnal ball-rolling dung beetles have been shown to use the celestial polarization pattern to set a straight path away from the food source where both predators and competitors may aggregate ^7,8^. In this case, dung beetles show the tendency to maintain the same heading over repeated trials ^6^. The tendency of other walking insects to set and maintain heading choices under a linearly polarized stimulus remains less well characterized. Although spontaneous behavioral responses to rotating polarization filters (polarotaxis) were demonstrated for crickets and flies when walking on air-suspended balls under laboratory settings ^9–11^, clear characterizations of angular heading choices are missing for these experiments. Similarly, behavioral data for flying insects (other than honeybees), especially when using virtual flight arenas remains relatively scarce ^1,2^. Oriented flights of suspended monarch butterflies under a polarized stimulus have been demonstrated, yet its ethological significance remains somewhat controversial, due to conflicting reports ^12–14^. Probably the most valuable recent progress comes from the fly *Drosophila melanogaster*: spontaneous responses of flying *Drosophila* to linearly polarized light using virtual flight arenas have been demonstrated, both under the natural sky, as well as using an artificial stimulus generated in the laboratory using commercially available polarization filters^15^,^16–18^. Most importantly, flies were shown to choose arbitrary angular headings with respect to the orientation of the e-vector of the polarized stimulus and showed the tendency to maintain this navigational decision over several minutes, even when the stimulus presentation was perturbed for several minutes ^18^. Nevertheless, fairly little is known about the navigational capabilities of free-living fruit flies (reviewed in ^19^). Catch-and-release experiments from a fixed point in the desert suggested that *Drosophila* (*melanogaster* and *pseudoobscura*) disperse into all directions equally and are able to keep straight headings over extended periods of time, while flying in environments which provide few visual landmarks ^20,21^. For a better quantitative understanding of the mechanisms underlying such processes, skylight navigation experiments using virtual flight arenas therefore serve as an attractive platform for the study of the navigation skills of wild type insects, thereby providing the platform for testing transgenic specimens harboring well-defined circuit perturbations ^22,23^.

The retinal basis of celestial polarization vision across insects is well understood: in virtually all cases, specialized ommatidia located in the ‘dorsal rim area’ (DRA) of the adult eye are morphologically and molecularly specialized for this task ^24^. In flies, DRA inner photoreceptors R7 and R8 express the same UV Rhodopsin Rh3 which is localized within untwisted light-sensing rhabdomeres, resulting in high polarization sensitivity ^22,25–32^. Since rhabdomeres of DRA R7 and R8 of a given ommatidium are oriented orthogonally to each other, these two cells form an opponent analyzer pair ^22,30^. Like in other insects, analyzer directions of DRA ommatidia change gradually along the DRA, forming a ‘fan-shaped array’ of polarization detectors ^33^. Due to the monochromatic Rhodopsin expression in DRA R7 and R8, navigational decisions of *Drosophila* in response to linearly polarized light should be limited to the UV range of the spectrum ^22,24^, whereas light of longer wavelengths should not elicit orientation responses to this stimulus. Interestingly, different insect species express blue-sensitive Rhodopsins in their polarization-sensitive DRA photoreceptors (crickets, locusts) ^34,35^, and in some cases green-sensitive Rhodopsins were reported (cock chafers) ^36^. Although these different Rhodopsin choices seem to reflect adaptations to different ecological niches, the exact ethological reason for these differences remain incompletely understood ^37,38^. Increasing evidence also points towards many insects (including flies) being capable to detect linearly polarized light through a DRA-independent channel (reviewed in ^39^). Experiments from Drosophila have shown that these polarotactic behaviors are not UV-specific, since behavioral responses can be elicited using polarized green light presented to the ventral half of the retina ^16,22^. Although incompletely understood, these behaviors could be indicative of a so-far poorly understood system in which retinal detectors are used to detect linearly polarized reflections. Such reflections could be used by insects to seek out or avoid water surfaces, evaluate oviposition sites, or even detect prey ^2,39^.

Previously, we have designed new and modular assays for studying visual navigation of single, flying flies in easy-to-build virtual flight arenas (for a detailed description, see https://doi.org/10.1101/527945 and www.flygen.org/skylight-navigation). Using this ‘open-loop’ setup, we now tested heading decisions of individual flies flying under a slowly rotating polarization filter. In agreement with previous studies, we find that flies initially choose a heading with respect to the orientation of the incident polarized light that varies between individuals and shows no preference for certain headings over the entire population tested (arbitrary headings). In the configuration used in our new assays, the rotation of the polarization filter therefore forces the fly to constantly adjust its heading in order to hold its original heading decision constant. By quantifying the fly’s ability to adjust its heading relative to the changing e-vector over time we show that the behavioral performance varies greatly within a population, yet a similar behavior is never observed under unpolarized UV light, or linearly polarized green light. Importantly, flies show the tendency to maintain this heading over several minutes: we show that flies that perform well in following the e-vector within a 5-minute experiment show a high tendency to choose a similar heading in a second experiment, even when interrupted by a 5-minute interval of unpolarized light. These experiments underscore the usefulness of the experimental setups presented here and serve as an ‘open source’ platform for the development of new assays optimized for different visual behaviors, in flies as well as other species of flying insects.

## Results

The aim of this study was a quantitative analysis of heading choices recorded from single flies flying under a slowly rotating polarization filter. We reasoned that flies that commit to a specific heading angle with respect to the incident angle of polarization would show a tendency to hold this angle constant and therefore correct for the slow rotational drift of the stimulus.

### New virtual flight arenas for studying skylight navigation in flies

In order to study heading choices of flies flying under linearly polarized light, we used a slightly modified version of the setup previously described by Mathejczyk and Wernet (https://doi.org/10.1101/527945). In short, by combining 3D printed mechanical parts with off-the-shelf hardware and computer vision methods we were able to quantify the behavioral responses of flying Drosophila (glued to a metal pin) to a constantly rotating e-vector of linearly polarized light presented dorsally (Figure 1A). This setup allowed individual magnetically tethered flies to rotate around their yaw axis, while keeping the flies’ position relative to the dorsal light stimulus constant. The polarization state of the dorsally presented stimulus was altered by shining light through a switchable filter ‘sandwich’ consisting of diffuser paper and a linear polarizer with either the polarizer or the paper facing the fly ^10,11,22^. This allowed for switching between two experimental conditions in which the degree of polarization was either 0% (unpolarized) and almost 100% linearly polarized while keeping light intensity between trials constant (Figure 1B). Due to the placement and size of the upper magnet (for holding the fly in place), the stimulus extended over a 28° wide concentric ring in the flies’ dorsal field of view (Figure 1C). By filming the flies from below and extracting their body axis angles over time using image processing their heading choices in response to the rotating e-vector were quantified (Figure 1D).

**Figure 1:**
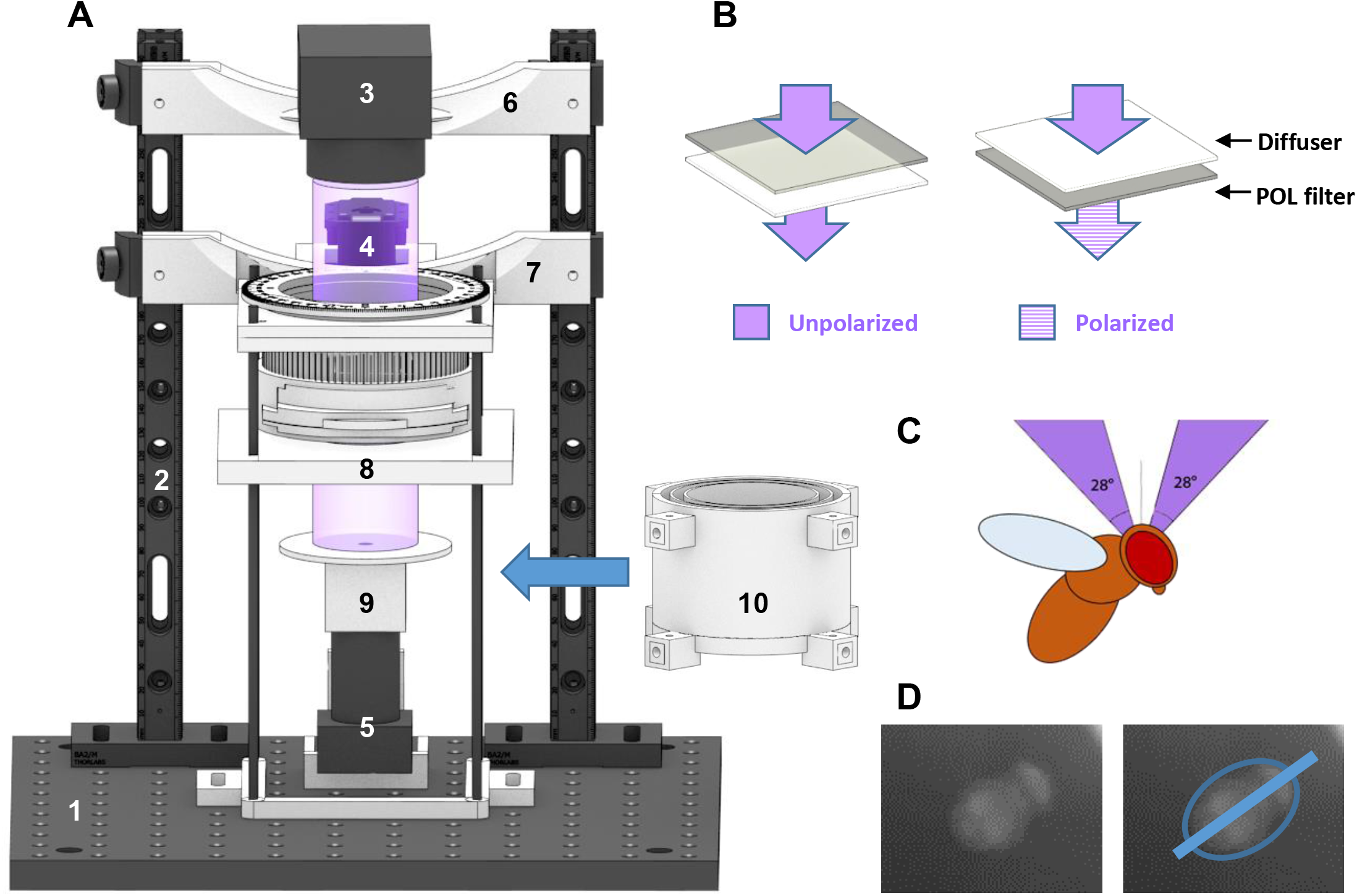
A new modular assay for studying skylight navigation in individual flying flies. **A.** Schematic drawing of a fully assembled virtual flight arena for the quantitative study of skylight navigation in flying *Drosophila*. Commercially available parts include: (1) a 30cmx30cm double density optical breadboard (Thorlabs); (2) compatible metal beams (Thorlabs); (3) swappable, magnetically attached high-power LED light sources with collimated optics (Mightex); (4) a robotics-grade servo motor for rotating the polarization filter (Dynamixel MX-28T, Robotis), positioned behind the light path (when viewed from the front); (5) an infrared camera shielded from the fly’s view, filming its body axis (Firefly MV, Point Grey). Custom-designed 3D-printed parts (see supplemental materials for printing instructions) include: (6) attachment to the vertical bars; (7) filter holder including gear system (see https://doi.org/10.1101/527945 and www.flygen.org/skylight-navigation for details); (8) horizontal platform holding the UV fused silica plate and top magnet for attaching the magnetotethered fly; (9) infrared illumination (enclosed LED’s) and bottom plate with ring magnet, mounted onto the infrared camera; (10) backlit cylinder for reducing linearly polarized reflection artifacts, which can be lifted all the way to the horizontal platform, when flies are tethered. **B.** Two polarization filter / diffuser orientations can be chosen within the filter holder: Unpolarized (diffused; left), or polarized (right). See Figure 3B for polarimetric characterization. **C.** Drawing of the fly’s field of view inside the apparatus. **D.** Camera image for the extraction of the fly’s body axis (shown in blue).

### Flying *Drosophila* follow a slowly rotating e-vector at an arbitrary angular distance

In order to achieve a more robust quantification of behavioral responses compared to rapid changes in e-vector orientation ^17^ (see https://doi.org/10.1101/527945), we introduced a new stimulus: flies flying within the virtual flight arena were presented a linearly polarized stimulus rotating slowly with constant angular velocity (~ 6°/s). In these experiments the 5-minute recording session per trial was split up into 30 × 10s windows. For each of these 30 windows the mean angular velocity of each fly was then calculated. If the difference between this angular velocity and the filter’s angular velocity was smaller than 3°/s, the particular time window was categorized as polarotactic behavior (areas shaded blue). For each of these periods the fly’s chosen heading was then calculated as the mean angular difference between the fly’s body axis and the incident e-vector (blue bar plots). A representative fly (Figure 2A) flying under a constantly rotating e-vector adjusted its heading in about one third of the recorded 10 sec time windows (10/30). The number and length of observed interruptions without polarotaxis varied from fly to fly, resulting in a wide spread of behavioral performance quality (as defined by number of polarotactic 10 sec time windows) when integrating over the entire 5 minutes tested. Importantly, the calculated preferred heading of a given fly falls within a narrow angular range when compared across polarotactic periods (Fly in Figure 2A: mean heading 63.2°, SD=12°), despite interspersed periods of non-polarotactic behavior. This indicates that flies attempt to keep a preferred heading with respect to the celestial e-vector pattern over short periods of time. As expected, virtually no polarotactic periods were detected when the same fly flew under unpolarized UV light (UVunpol), but otherwise unchanged conditions (Figure 2B). Similarly, when the fly was flying under linearly polarized green light (PolGreen), virtually no polarotaxis was detected (Figure 2C). However, upon presenting the fly re-polarized UV light again (PolUV2), polarotactic time periods were restored, in some cases even more pronounced than in the first UV trial (Figure 2D). Pooled data from all tested flies flying under a constantly rotating filter under different lighting conditions (PolUV1, UVunpol, PolGreen, PolUV2) reveals that polarotaxis occurs exclusively when using a linearly polarized UV stimulus (Figure 3A). Flies flying under a slowly rotating linearly polarized UV stimulus spent significantly more time following the e-vector, compared to flying under unpolarized UV or polarized green light, respectively. Interestingly, flies that underwent a second flight under re-polarized UV light spent even more time following the e-vector. Finally, analysis of behavioral responses of female and male flies reveal no significant differences between genders in the first as well as in a second trial under polarized UV light (PolUV1 and PolUV2).

**Figure 2:**
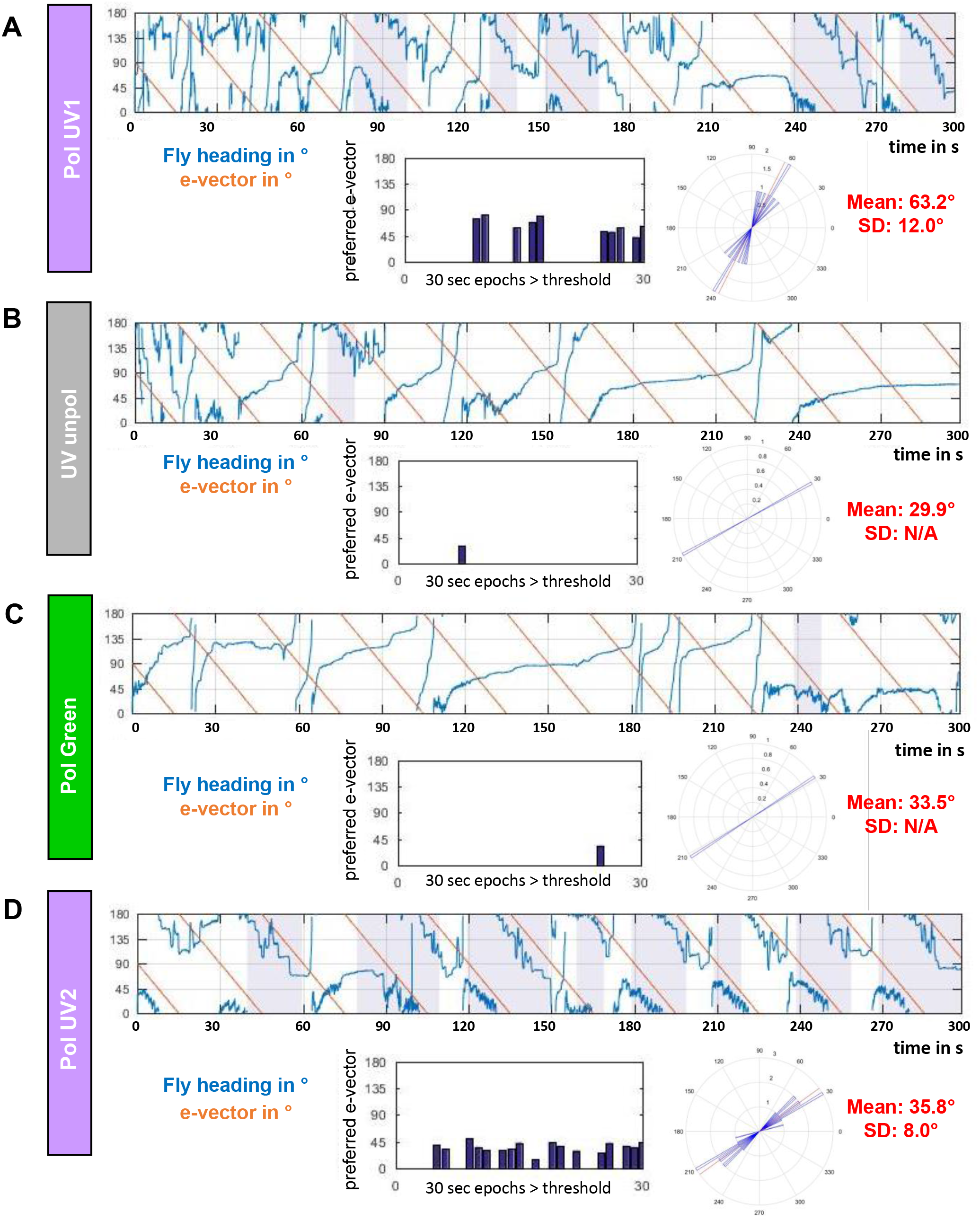
Flying *Drosophila* follow a slowly rotating e-vector at an arbitrary angle. **A.** Left: Flight heading (blue line) of a single fly orienting to a slowly rotating linearly polarized UV stimulus (PolUV1; orange line). 30 s intervals with above threshold polarotactic behavior (see material and methods) are shown in grey. Right: plot of above threshold intervals and circular plot of heading chosen by the animal. **B.** Same analysis as above, using an unpolarized UV stimulus (UVunpol); same fly. **C.** Same analysis as above, using a polarized green stimulus (PolGreen); same fly. **D.** Same analysis as above, using a re-polarized UV stimulus (PolUV2); same fly.

**Figure 3:**
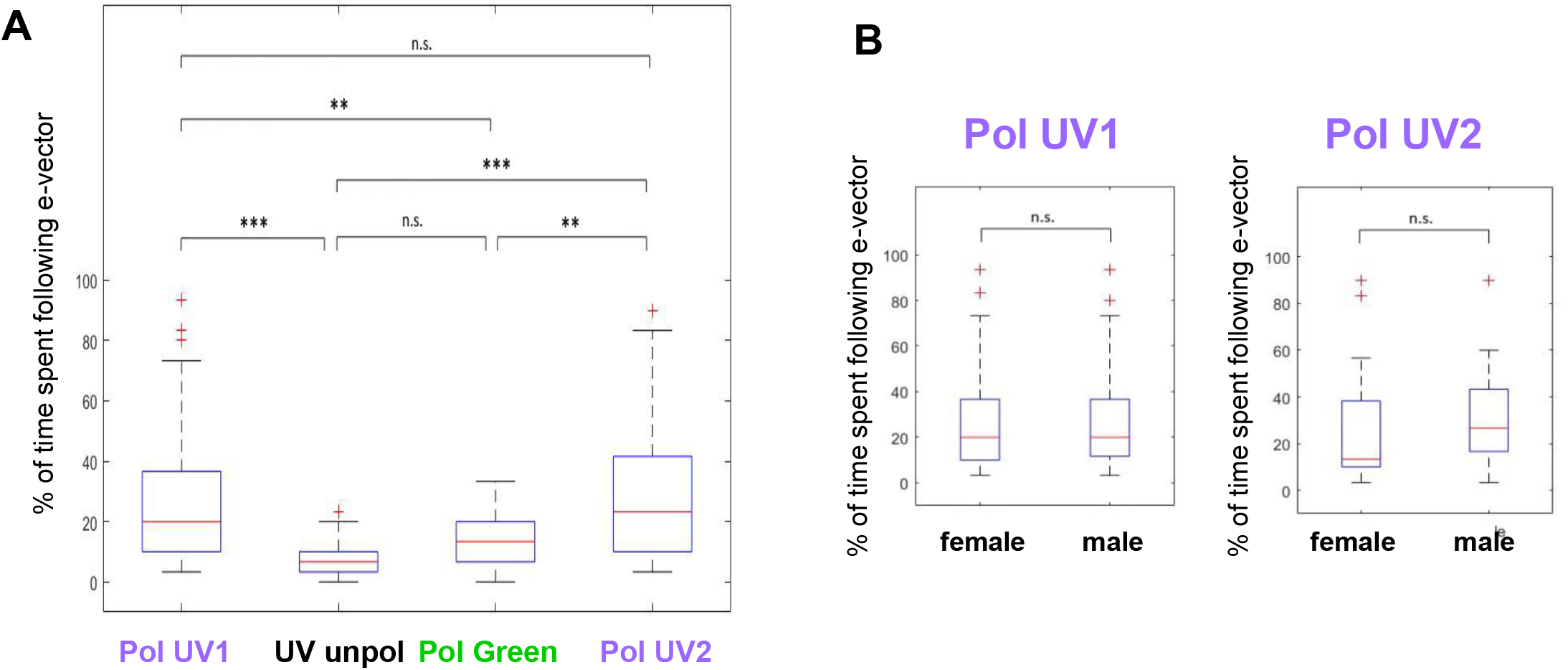
Variation of heading choices across individuals tested. **A.** Summary plot of % time spent following the slowly rotating e-vector, comparing the four conditions from above. Bonferroni-corrected two-tailed Mann-Whitney U test, *p<0.05, **p<0.01, ***p<0.001. N (left to right) = 66, 22, 38, 43. **B.** Direct comparison of male versus female polarotaxis from PolUV1 and PolUV2 experiments reveals no significant difference. Two-tailed Mann-Whitney U test. N (left to right) = f (30), m (36), f (19), m (24).

### Chosen headings are arbitrary, while behavioral performance varies between flies

By comparing the behavior across many individuals (N = 66) we investigated the spread of preferred headings when single flies were flying under a slowly rotating polarization filter. The goal was to investigate whether, in this particular kind of virtual flight arena, certain headings are naturally preferred or avoided, or whether the choice of preferred heading is arbitrary and therefore different between individual flies. The strategy is exemplified by the direct comparison of four representative traces of individual flies in response to a slowly rotating polarized UV stimulus (Figure 4). Quality of behavioral performance (polarotaxis intervals) and preferred heading were quantified as described above. It appears that in these trials each fly choses a different preferred heading with respect to the incident angle of polarization (circular plots). Taken together, the preferred heading angles of all tested flies during their first linearly polarized UV trial (PolUV1) were distributed over the whole angular range (Fig. 5A,C). Importantly, although the quality of behavioral performance (number of polarotaxis intervals, i.e. time spent following the rotating e-vector) varied greatly between individuals, it did not correlate with the angular heading choice of the animals (Fig. 6A). Hence, unlike previous studies on walking fly populations ^22^, no tendency for a fixed preferred angular heading choice was found across flying individuals.

**Figure 4:**
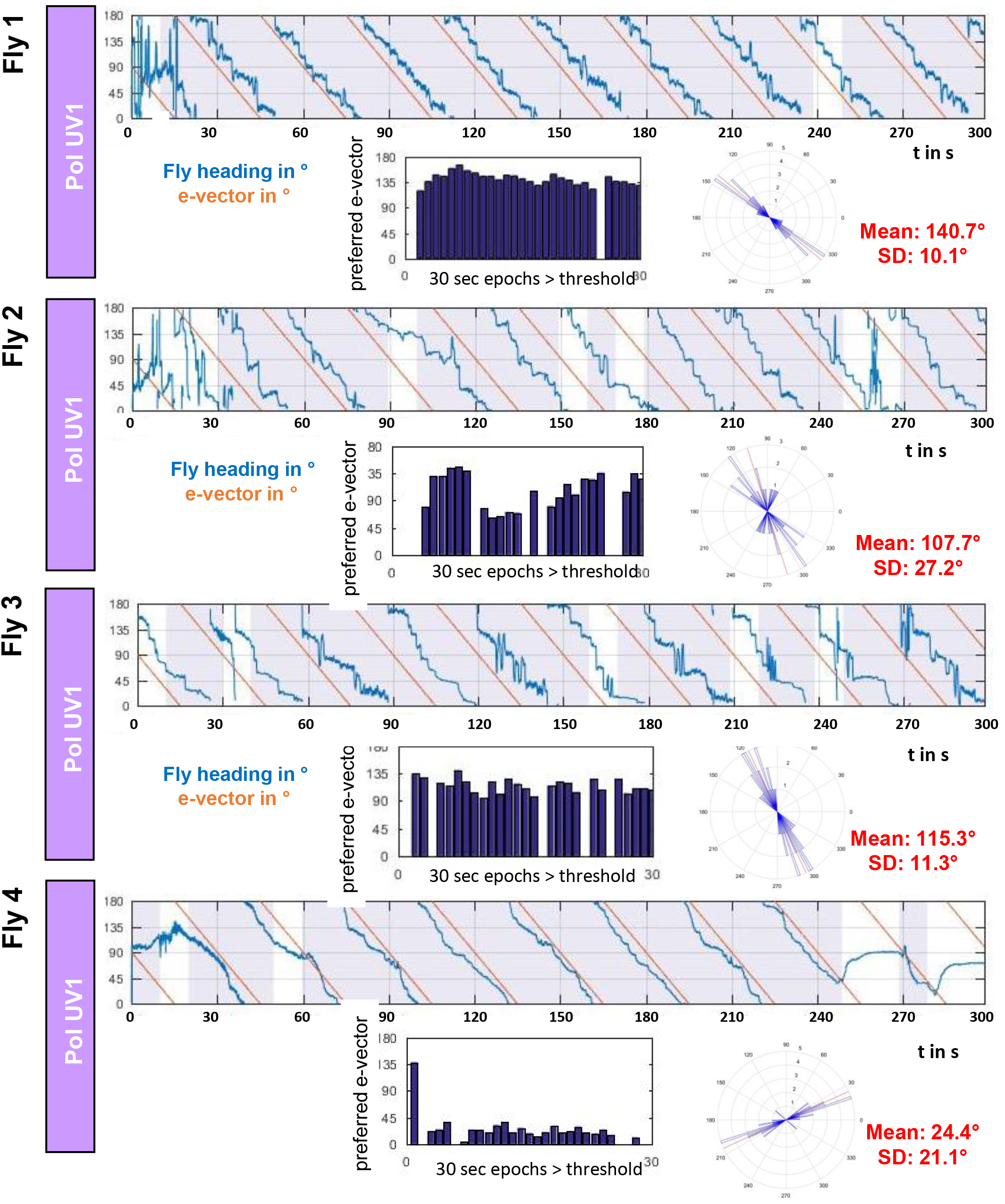
Chosen headings are arbitrary and different between flies. Flight heading (blue line) of four single flies orienting to a slowly rotating linearly polarized UV stimulus (PolUV1; orange line). 30 s intervals with above threshold polarotactic behavior (see material and methods) are shown in grey. Right: plot of above threshold intervals and circular plot of heading chosen by the animals.

**Figure 5:**
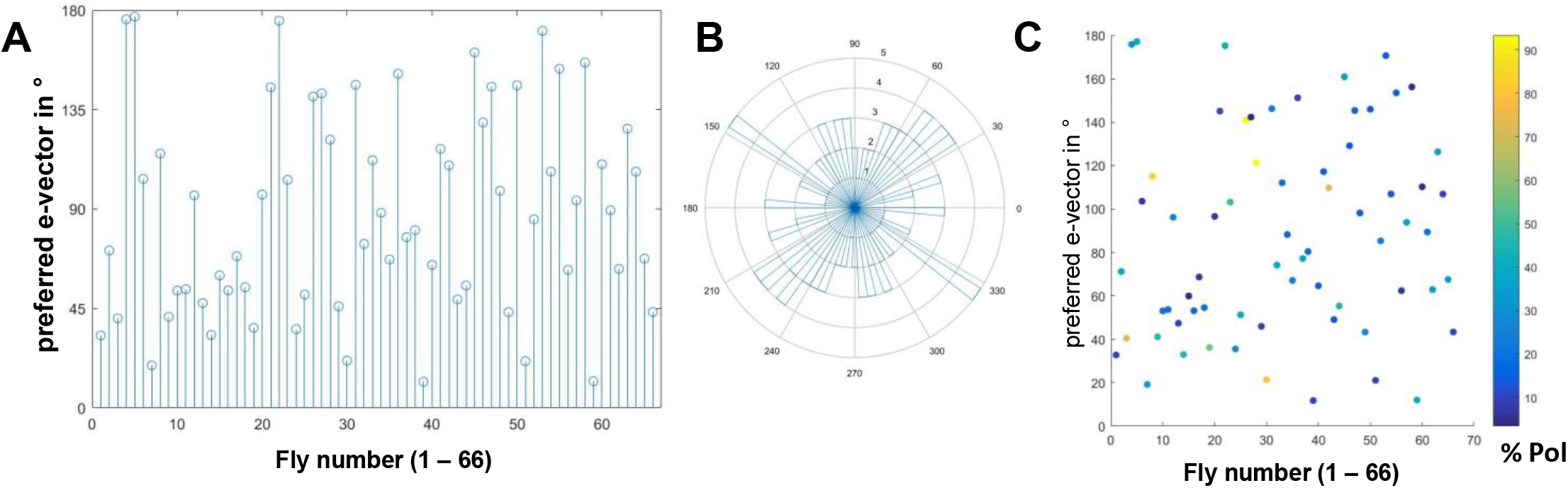
Variability and robustness of consecutive heading choices. **A.** Summary of heading angles (angular difference to the polarization filter) chosen by 66 individual flies, under the PolUV1 stimulus. **B.** Same data plotted on circular coordinates reveals a wide distribution. **C.** Same data as in B where behavioral performance (% of time following the e-vector) is represented in false color.

**Figure 6:**
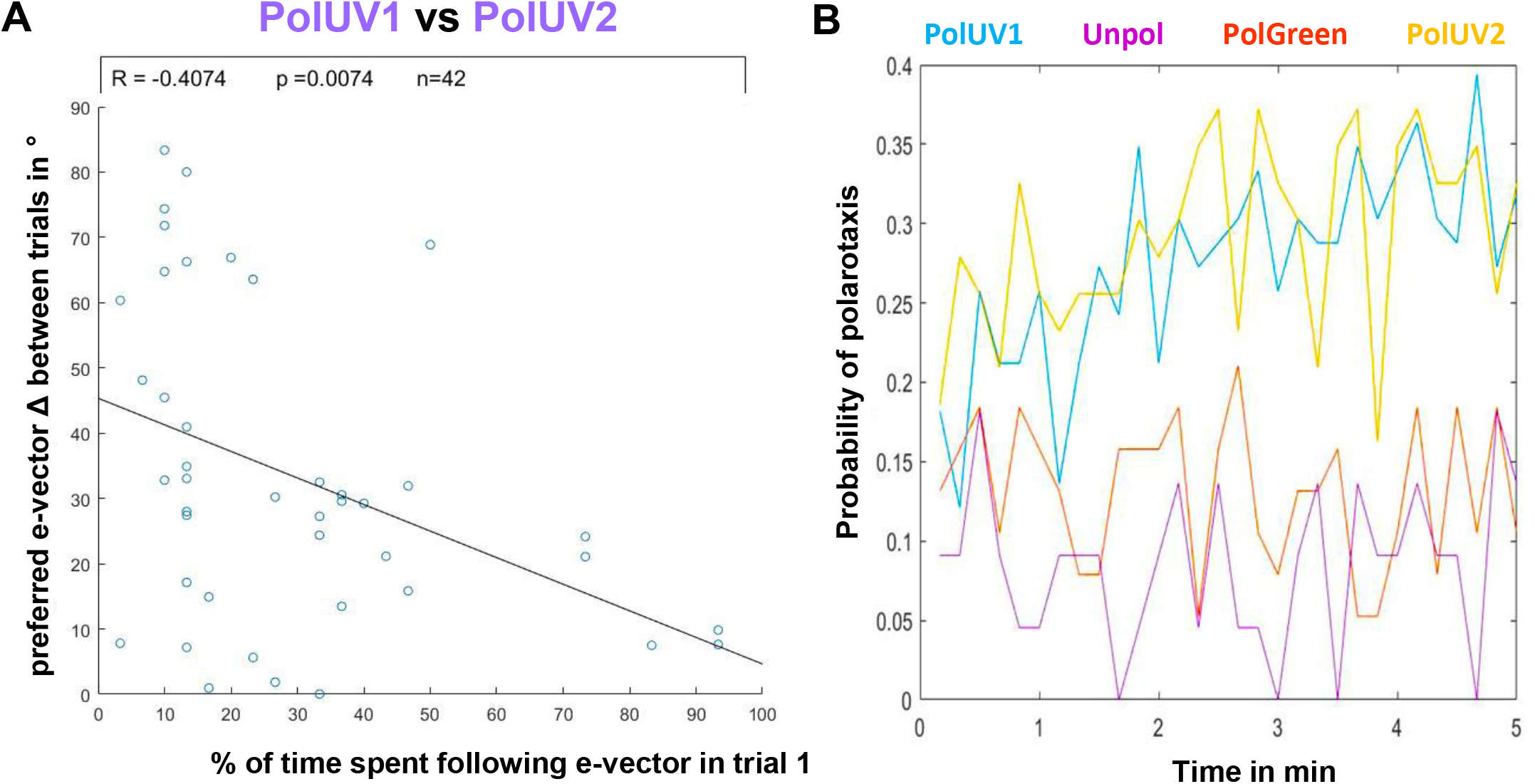
Arbitrarily chosen headings are maintained between trials. **A.** Plot depicting the angular difference between headings chosen in two consecutive trials interrupted by a period of unpolarized light (PolUV1 vs PolUV2) as a function of the quality of polarotaxis during the PolUV1 period (% of time spent following e-vector). B. The change of polarotaxis probability over the 5 min experimental time window, plotted for all 4 experimental conditions (PolUV1, UVunpol, PolGreen, and PolUV2).

### Arbitrarily chosen headings are maintained between trials

Finally, we tested whether the amount of time that the flies spent following the e-vector (number of polarotaxis intervals, i.e. quality of behavioral performance) within the first linearly polarized UV trial (PolUV1) correlates with the tendency of the flies to choose a similar preferred heading in a second consecutive trial (PolUV2) that was separated from the first by an interruption (5 min of UVunpol). We found that the better the flies’ performance within the first trial (more time spent following the e-vector in PolUV1), the higher the likelihood of them choosing a similar heading in the second trial (Figure 6A). During both PolUV1 and PolUV2 intervals, the probability of tested flies for following the rotating e-vector increased during the 5 min trial (Figure 6B). In contrast, the overall lower polarotactic values obtained in control conditions (UVunpol and PolGreen) showed no similar increase over time.

## Discussion

Navigating insects rely on the detection and integration of a wide variety of visual cues, like celestial bodies (sun, moon, milky way), intensity gradients, and chromatic gradients ^1^. In addition, the celestial pattern of linearly polarized light serves as an attractive orientation cue that many insects use ^2,40,41^. Spontaneous behavioral responses of both walking and flying *Drosophila* to linearly polarized light (‘polarotaxis’) have been demonstrated in the past, using both population assays, as well as single fly assays^16–18,22,33,42,43^. In all these experiments, much care was given to the control and avoidance of intensity artifacts that can result in behavioral decisions that are in fact independent of the linearly polarized component of the stimulus (reviewed in^15^). However, some of the successful solutions presented in the past included components whose reproduction required considerable engineering skills and were quite costly. The virtual flight arenas used here have been designed with the dual goal of providing relatively cheap, robust setups that can easily be assembled, while at the same time minimizing intensity/reflection artifacts. The codes, templates and building instructions for the virtual flight arenas are freely available for download to anyone (for a detailed description, see https://doi.org/10.1101/527945 and www.flygen.org/skylight-navigation). Due to their modular design, the use of our new virtual flight assays is in no way limited to skylight navigation or the study of polarized light vision. With few simple modifications they could be modified for studying behavioral responses to moving stimuli ^44–47^, shapes ^48^, colors ^49^, or celestial bodies ^23^. Similarly, the setups can easily be modified to house a spherical treadmill, for studying the visual behavior in walking flies. Finally, application is in no way limited to just *Drosophila* or other flying insects. We hope that the assays used here (and to be published elsewhere in great detail, including assembly instructions) can serve as a platform from which many other assays optimized for many other species could evolve from.

Using these new virtual flight arenas, we show that individual flies choose arbitrary headings under a linearly polarized stimulus, and when summed over all individuals tested, all chosen headings appear to spread randomly. This finding is in good agreement with recently published studies, although these were using a rather different kind of stimulation system^18^. In further agreement with these past studies, we also find that flies show a clear tendency to keep their chosen heading over several minutes, which indicates that any given fly attempts to maintain its chosen heading. Given the considerable gap in knowledge about the ethology of *Drosophila*, these data provide further support for a potential role of polarization vision in guiding long-range navigation behaviors that have been reported for flies ^19–21^. In contrast, we show that single flies flying under a linearly polarized green stimulus displayed no comparable polarotaxis, which was to be expected due to the fact that polarization-sensitive R7 and R8 photoreceptors in the dorsal rim area (DRA) ^24^ of the fly eye both express the UV-sensitive Rhodopsin Rh3^27–29^. Nevertheless, behavioral responses to linearly polarized stimuli with longer wavelengths have also been reported in the past ^16,22^, especially when presented ventrally, and the retinal detectors responsible are not known ^22,39^.

Our experiments show that well-performing flies show a clear tendency to maintain their chosen heading, even when interrupted by a period of unpolarized stimulation. These data again provide independent support for previous studies ^18^ and reinforce the idea that a generalist fly like *Drosophila melanogaster* is indeed capable of using skylight polarization for maintaining a chosen course over longer times, which is crucial for achieving more complex navigational tasks ^19–21^. Like previous studies, we aimed at quantifying the quality of behavioral responses, since we expected that behavioral performance of individual flies to be greatly variable due to the strong influence of environmental conditions as well as internal states of the animal(s). For this study, we introduced a simple new stimulus, where flies are suspended under a slowly rotating polarization filter under ‘open loop’ conditions. Quantifying the quality of a behavioral response by chopping any given 5-minute experiment into 30×10 sec polarotactic periods serves as an attractive new strategy for producing statistically significant data in a reasonable amount of time. Using this method, our experiments indeed revealed that behavioral performance is variable across all individuals tested. Even after tight control of food quality, rearing conditions, temperature, and humidity, the flies’ cooperation in these experiments remains unpredictable. How much this variability could depend on the fly’s motivational state or navigational decision making remains to be investigated. Interestingly, even within a given 5 min recording, flies do not necessarily follow the rotating e-vector permanently, but may transition into and out of polarotactic periods (Figure 2A). This demonstrates the usefulness of this experimental setup for further studies on the dynamics and modulation of polarotactic behavior (and potentially underlying decision making processes), for instance in response to different internal states. Finally, our experiments reveal no significant differences in the behavioral performance of male versus female flies. This was to be expected since catch-and-release experiments did not reveal any sex differences in *Drosophila’s* tendency to disperse ^20,21^. Furthermore, the size and structure of DRA ommatidia does not differ in a systematic way, between sexes. Although male-specific, Fruitless-expressing neurons have been characterized in the Drosophila brain, none of them appear to be clustered in the dorsal periphery of the visual system ^50,51^.

Many insect species use the celestial polarization pattern in conjunction with other visual stimuli like celestial bodies, intensity gradients, chromatic gradients, and landmarks ^1^. The hierarchy in which these stimuli are combined might differ between species as well as depending on context. One recent study reported that single *Drosophila* flying in a virtual flight arena are also able to use an artificially generated celestial body (the sun) as a reference to choose a heading (menotaxis) ^23^, a behavior that requires ‘compass neurons’ in a central brain region known as the central complex ^52^. This function is therefore in good agreement with physiological properties described for these neurons in locusts ^53^. Classic data from larger insects ^5^, as well as more recent studies from *Drosophila* ^54^ are beginning to elucidate the neural circuitry of the ‘compass pathway’, along which menotactic and polarotactic information are being integrated by the insect brain, resulting in time-compensated compass information in the central complex ^5,55^. Despite important similarities, it remains unclear whether flies use their anatomical compass pathway for performing exactly the same computations for skylight navigation ^23,54,56–60^. The experiments presented here therefore serve as an important new platform for the efficient combination of *Drosophila* molecular genetic tools for the cell-type specific manipulation of neuronal function with quantitative behavior assays for testing skylight navigation.

## Methods

### Fly rearing

Wild type Oregon R flies (isogenized) were reared at 25°C and 60% relative humidity on standard cornmeal agarose food under a 12h-light/12h-dark cycle. Care was taken to keep population densities low within fly vials by flipping flies on a daily basis.

### Fly preparation

Experiments were performed at 25°C and 50% relative humidity during the flies’ evening activity peaks up until one hour after the light period within the respective rearing incubators would have ended. Flies were glued to 10mm long, 100μm diameter steel pins (ENTO SPHINX s.r.o., Czech Republic), so that when positioned vertically, they held the flies at a natural flying angle (about 60° from horizontal) and were allowed to recover for at least 20 minutes from the gluing procedure before being tested. To prevent the flies from flying during the recovery phase, small pieces of Kimwipes were transferred to their tarsi. Initial flight behavior was triggered by inducing a little air puff towards the fly from below. This was also quickly done when flies stopped flying during the experiments, but not more than 3 times per experiment without excluding those flies from data analysis.

### Flight simulator setup

#### Virtual flight arenas

A detailed description of the virtual flight arenas used in this study including building instructions and the codes necessary for their operation are described in detail elsewhere (https://doi.org/10.1101/527945 and www.flygen.org/skylight-navigation). In short, tethered flies were placed between two magnets, allowing them to rotate freely around their yaw axis, while the magnetic field of the magnets kept the steel pins vertical. A sapphire bearing at the bottom side of the upper magnet minimized friction and kept the steel pins in place. Flies were filmed from below with 60hz (Firefly MV, Point Grey) under near-infrared illumination.

#### Stimulus delivery

Above the fly a switchable cassette holding a 50mm × 50mm sheet linear polarizer (OUV5050, Knight Optical, UK) and 13 layers of thin, non-fluorescent diffuser paper (80g/sqm, Max Bringman KG) was inserted into a motorized rotatable cassette holder. Light from a collimated UV or green LED (365nm: LCS-0365-13-B, 530nm: LCS-0530-15-B, Mightex) coming from above and passing through the filter cassette was polarized (pol filter at bottom) or depolarized (diffuser paper at bottom) depending on the orientation of the filter cassette and presented to the dorsal part of the flies’ eyes. Using a spectrometer (Flame, Ocean Optics) the intensity of the two LEDs was set approximately isoquantaly at 2×10^12^ photons/s/cm^2^. By rotating the cassette holder, it was possible to precisely control the angle of the polarized stimulus’ e-vector. The recording protocol consisted of rotating the E-vector with constant angular velocity (5.97 deg/s) for 5 minutes while synchronously recording the flies’ behavior.

#### Extraction of flight heading

The heading of each fly in each acquired video frame was extracted using a custom-written macro script for the open-source software Fiji ^61^. In short, it binarizes each video and fits an ellipse around the fly’s body to extract its heading in a range from 0° to 180°, in accordance with the directional ambiguity of the presented e-vector. The Fiji tracking results were analyzed using MatLab. Circular statistics were used. In short, a fly’s heading changes were quantified as polarotactic behavior if the mean difference between the fly’s angular velocity and the e-vector’s angular velocity was smaller than 3°/s for a given 10s time window. By calculating the fly’s heading relative to the e-vector during such polarotactic episodes it was possible to calculate a measure of the ‘preferred e-vector’ for each fly. The custom Fiji- and Matlab-scripts used in this study, as well as setup building instructions and documentation is freely available under www.flygen.org/skylight-navigation.

## Supporting information

Mathejczyk and Wernet Supplemental Data BioRxiv

## Acknowledgements

The authors thank Leigh Moss, Tanja Heinloth, Lena Naber and Nurelhoda Abdel Muti for experimental support, Dr. Gerit Linneweber for statistical advice, as well as all members of the Wernet and Hiesinger groups for their input. This work was supported by the Deutsche Forschungsgemeinschaft through grants WE 5761/2-1 and SFB958 (Teilprojekt A23), through AFOSR grant FA9550-19-1-7005, through the Berlin Excellency Cluster NeuroCure, with support from the Fachbereich Biologie, Chemie & Pharmazie of the Freie Universität Berlin, as well as the Division of Neurobiology at Freie Universität Berlin (support of FU Berlin and the National Institute of Health to Robin Hiesinger).

## Author contributions

TFM and MFW planned the experiments. TFM built the assay and performed all experiments. TFM and MFW designed the figures. MFW wrote the manuscript, MFW and TFM finalized the manuscript.

## Data availability

The datasets generated during and/or analysed during the current study are available from the corresponding author on reasonable request.

