## Supplementary material for "Heading choices of flying *Drosophila* under changing angles of polarized light": Mathejczyk and Wernet Supplemental Data BioRxiv

**Supplemental Material**

Supplemental Figure S1: Setup overview

**A.** Picture of a single flight simulator. (1) Collimated stimulus LED; (2) rotatable filter cassette holder; (3) filter cassette containing sheet polarizer and paper diffuser; (4) upper magnet; (5) movable cylinder containing white LEDs. **B.** Picture of the temperature- and humidity-controlled enclosure containing a humidifier (middle), a heating plate (back) and 2 flight simulators, one with a green (left) and one with a UV stimulus LED (right). **C.** Same as B, but with the white LEDs within the movable cylinder turned off for better visualization of the polarized stimulus light path. For a detailed description, see https://doi.org/10.1101/527945 and www.flygen.org/skylight-navigation.

Supplemental Figure S2: Test for behavioral impairment

Bar plots depicting the sum of rotational motion for all tested flies per 5 min trial (positive = clockwise, negative = counter-clockwise), showing that the flies‘ ability to rotate around the yaw axis is not impaired, for any of the experimental conditions tested.

Supplemental Figure S3: Flies are able to keep a chosen heading relative to the evector even after a 5min break

**A-C.** Scatter plots showing the chosen preferred e-vector over consecutive trials UVPol1/UVPol2, UVPol1/UVUnpol1 and UVPol1/GRPol1, respectively, for all tested flies. Shading of points indicates percentage of time each fly spent following the e-vector rotation in the first trial. Under polarized UV conditions flies that spent the most time following the e-vector rotation in the first trial are better at keeping this chosen angle in a second trial (indicated by clustering of dark points around the diagonal black line) compared to unpolarized UV and polarized green light, where this effect seems much weaker. **A‘-C‘.** In order to assess whether a preferred e-vector might be chosen randomly in a second trial, the mean angular difference between the preferred e-vectors chosen in two consecutive trials that was measured (black arrows) was compared to the distribution of the mean angular differences between the measured preferred e-vectors chosen in the first trial and sets of randomized e-vectors (shuffled 10000 times, blue distribution). Under polarized light the measured mean differences between the preferred e-vectors chosen in two consecutive trials show lower probabilities of occurring randomly compared to flies flying under unpolarized light. N = A/A‘:42; B/B‘:20; C/C‘:35.

### Supplemental Figure S1

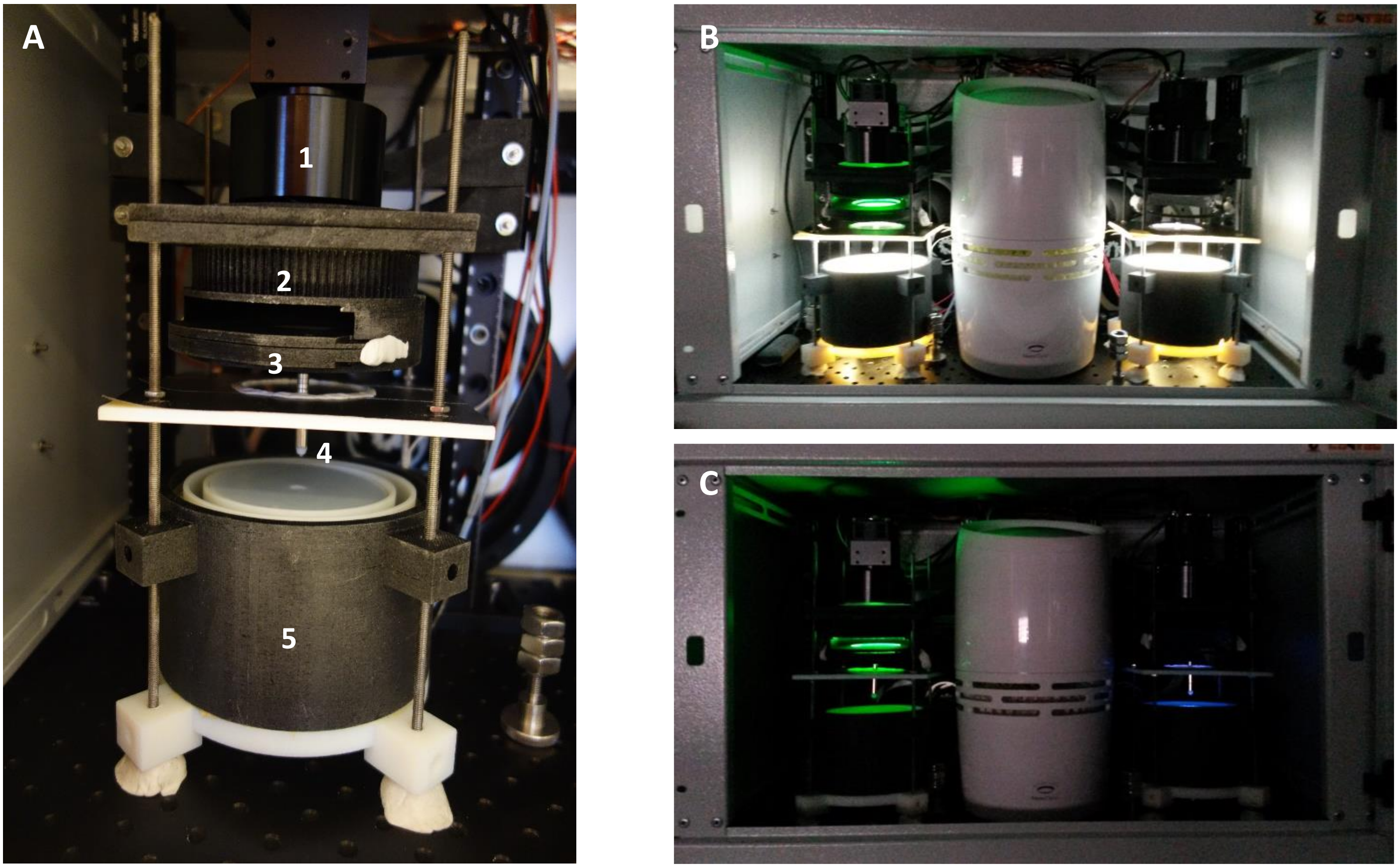

#### Supplemental Figure S2

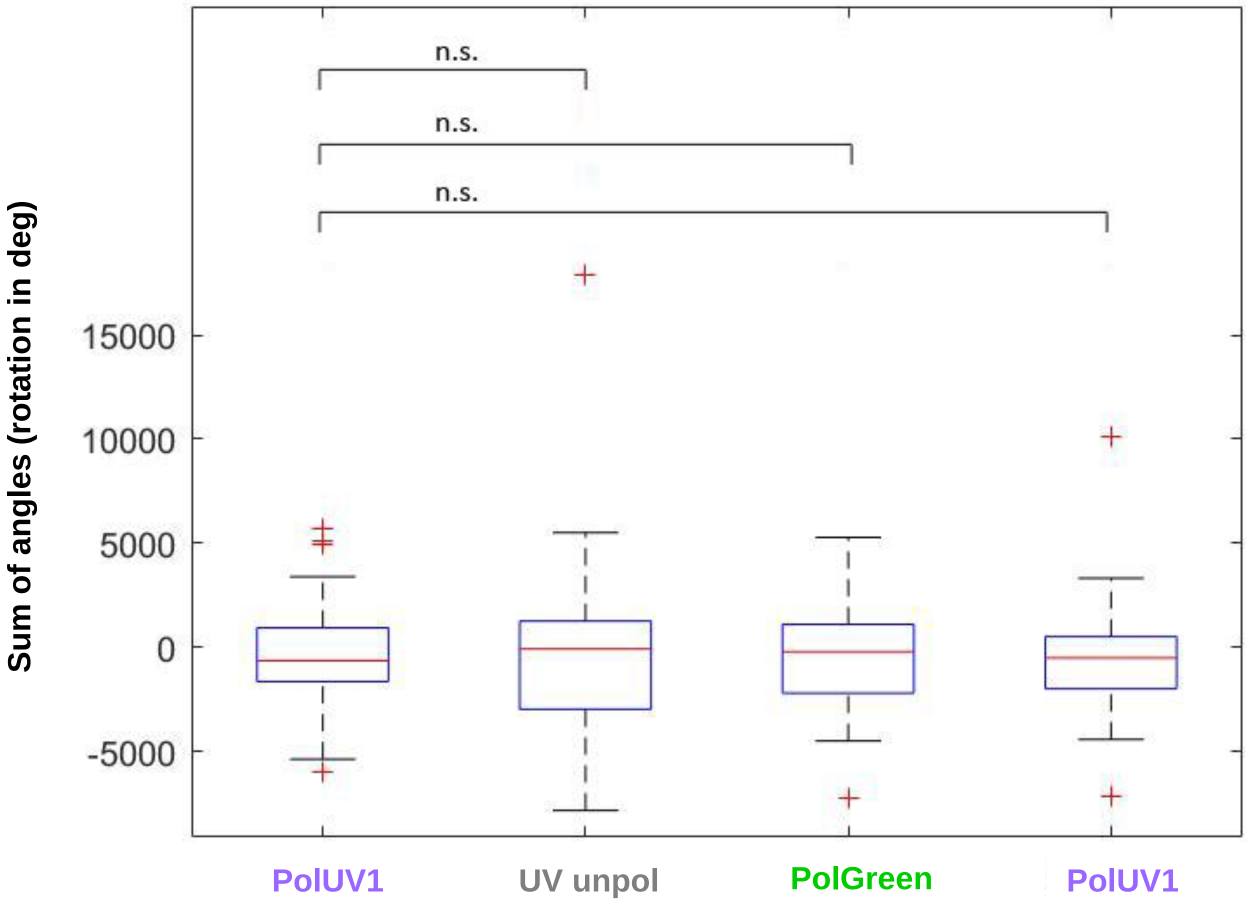

### Supplemental Figure S3

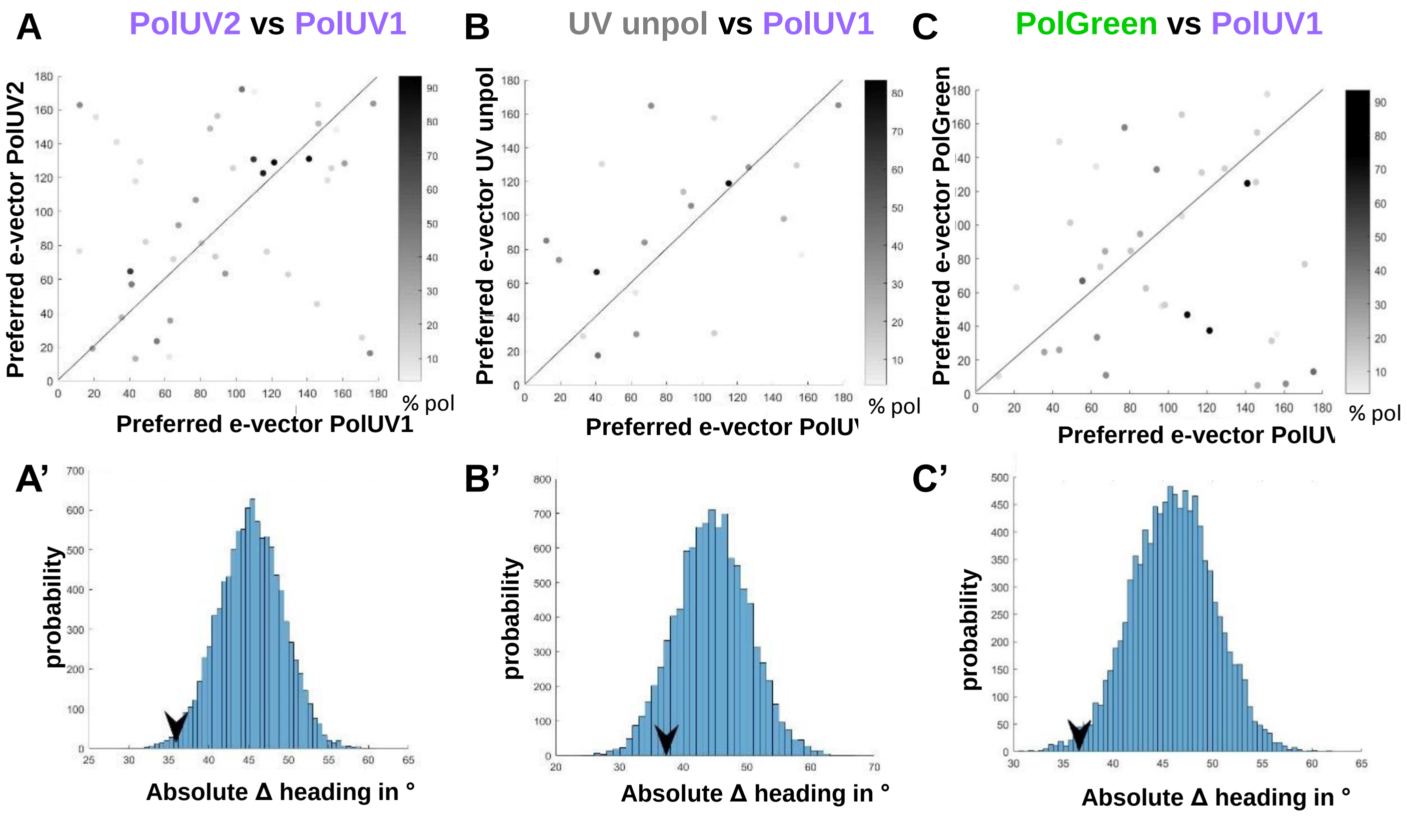
